# Tankyrase-mediated ADP-ribosylation is a novel regulator of TNF-induced death

**DOI:** 10.1101/2021.02.09.430424

**Authors:** Lin Liu, Jarrod J. Sandow, Deena M. Leslie Pedrioli, Natasha Silke, Zhaoqing Hu, Emma Morrish, Diep Chau, Tobias Kratina, Andrew J. Kueh, Michael O. Hottiger, Andrew I. Webb, Najoua Lalaoui, John Silke

## Abstract

Tumor necrosis factor (TNF) is an inflammatory cytokine that, upon binding to its receptor TNFR1, can drive cytokine production, cell survival, or cell death and is a major component of an organism’s anti-pathogen repetoire^1,2^. TNF stimulation leads to the formation of two distinct signalling complexes, a well-defined membrane bound complex (complex 1), and a less well characterised cytosolic death inducing complex (complex 2). Using mass spectrometry, we identified the ADP-ribosyltransferase, tankyrase-1 (TNKS1/TNKS/ARTD5/PARP5a) as a novel native complex 2 component. Following a TNF-induced death stimulus TNKS1 is recruited to complex 2, resulting in complex 2 poly(ADP-ribosyl)ation (PARylation). Tankyrase inhibitors sensitise cells to TNF-induced death, which is correlated with increased complex 2 assembly. Tankyrase-mediated PARylation promotes recruitment of the E3 ligase RNF146 and RNF146 deficiency or proteasome inhibition results in increased levels of complex 2, suggesting that RNF146 causes proteasomal degradation of complex 2. Several viruses express ADP-ribose binding macrodomain proteins, and expression of the SARS-CoV-2 or VEEV macrodomain markedly sensitises cells to TNF-induced death. This suggests that ADP-ribosylation serves as yet another mechanism to detect pathogenic interference of TNF signalling and retaliate with an inflammatory cell death.

---

Tumor necrosis factor (TNF)/TNFR1 signalling helps coordinate an anti-pathogen response by promoting transcriptional upregulation and secretion of other cytokines and inflammatory mediators ^3–17^. To counter this, pathogens have evolved mechanisms to disrupt signalling from the membrane bound complex 1 that nucleates around TNFR1 ^2,18^. This in turn has prompted an evolutionary arms race whereby disruption of the transcriptional response can provoke TNF-induced cell death via a secondary cytosolic complex 2, containing RIPK1, FADD and caspase-8^3,5,16,19–32^. Dysregulation of TNF signalling has been implicated in a diverse range of inflammatory and auto-immune diseases ^33–35^, stimulating research that has generated a detailed understanding of complex 1 and the TNF/TNFR1 transcriptional response. Compelling evidence showing that TNF-induced cell death is also pathogenic has stimulated the development of drugs to block the cell death response ^16,35–37^, but a correspondingly detailed insight into the composition and regulation of complex 2 is lacking.

## Tankyrase-1 is a novel component of TNFR1 complex 2

To identify TNFR1 complex 2 components, we generated and validated both N- and C-terminally 3x FLAG tagged murine caspase-8 constructs (**Extended Data Fig. 1a-c)**. These tagged constructs allowed us to immunoprecipitate caspase-8 with a number of controls that increase the chance of identifying true hits. Complex 2 formation was induced by treating cells with TNF (T), Smac-mimetic (S) to impair the transcriptional response and the pan-caspase inhibitor emricasan/IDN-6556 (I) to stabilise complex 2 ^38–40^. As expected, mass spectrometry analysis of the caspase-8 C3FLAG immunoprecipitate from TSI treated Mouse Dermal Fibroblasts (MDFs) revealed enrichment of known complex 2 components, including RIPK1, RIPK3, A20, TRADD and FADD (**Fig. 1a; Supplementary Data 1, sheet 1**). We also identified a previously unreported complex 2 protein, tankyrase-1 (TNKS/TNKS1/ARTD5/PARP5a) (**Fig. 1a; Supplementary Data 1, sheet 1**). TNKS1 is an ADP-ribosyltransferase of the ARTD family ^41,42^ (**Extended Data Fig. 1d**), and has not previously been implicated in regulating TNF-induced cell death. To explore the physiological significance of this finding we generated both N- and C-terminally 3x FLAG tagged caspase-8 (*Casp8^N3FLAG^* and *Casp8^C3FLAG^*) knock-in mice using CRISPR/Cas9 technology (**Extended Data Fig. 1e-f**). Bone marrow derived macrophages (BMDMs) and MDFs generated from heterozygote knock-in mice were treated with TSI and caspase-8 was immunoprecipitated ± FLAG peptide spiking. As expected, cleaved caspase-8, FADD and RIPK1 were immunoprecipitated together with caspase-8 upon TSI from both *Casp8^+/N3FLAG^* and *Casp8^+/C3FLAG^* cells although we precipitated slightly more of these proteins from *Casp8^+/C3FLAG^* cells (**Fig. 1b**, **Extended Data Fig. 1g**). Consistently we also observed higher levels of TNKS1 co-precipitating with caspase-8 C3FLAG **(Fig. 1b**, **Extended Data Fig. 1g**). In contrast, we did not observe PARP1/ARTD1, the most widely studied ARTD family member, co-precipitating with caspase-8 after TSI stimulation, suggesting that the association of TNKS1 with complex 2 was specific (**Extended Data Fig. 1h**).

**Fig. 1.**
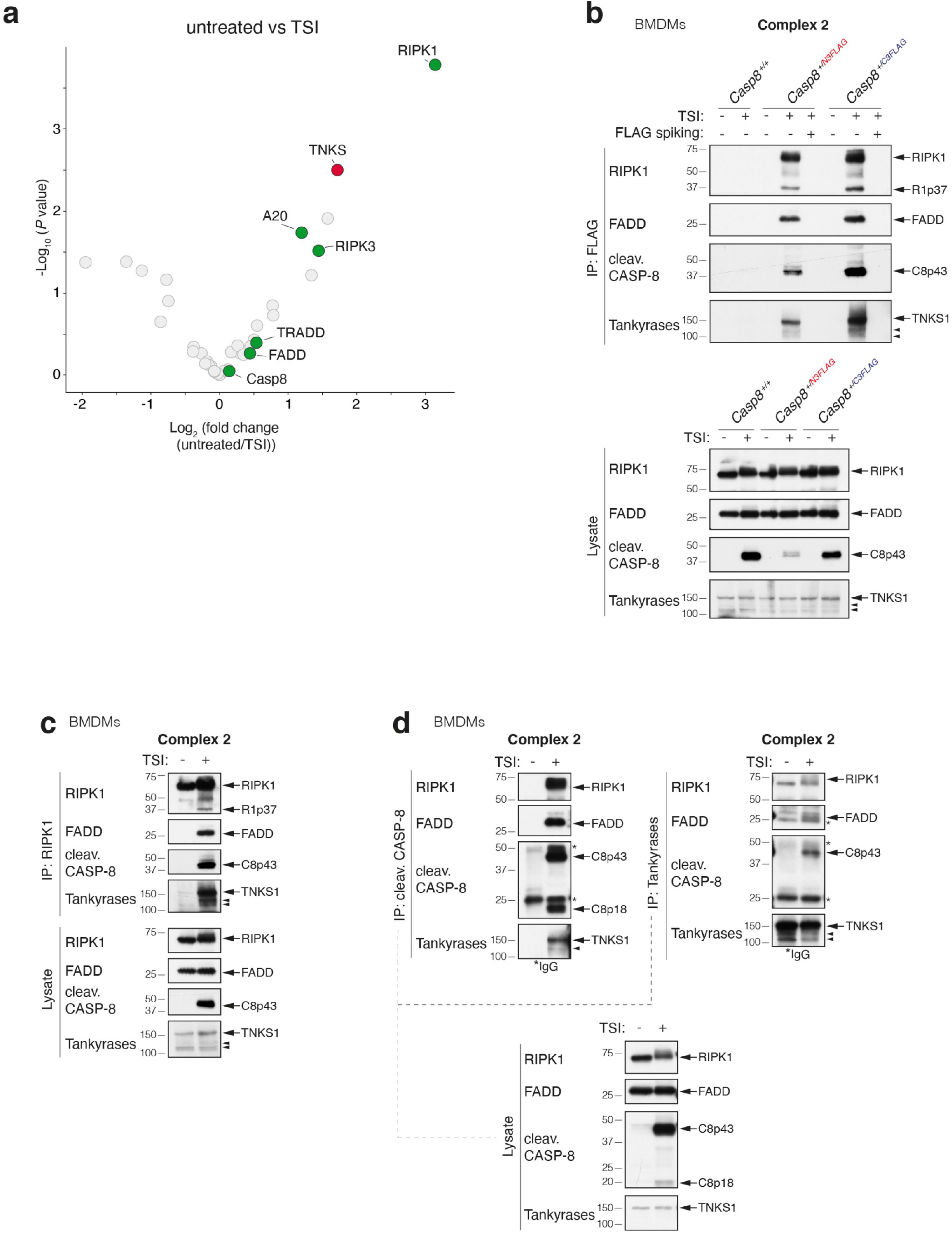
Tankyrase-1 is a novel interactor of native TNFR1 complex 2. **a,** Log2 fold change volcano plots of protein enrichment upon TSI stimulation in *Casp8^-/-^.Mlkl^-/-^* MDFs expressing caspase-8 C3FLAG compared to the untreated control. Proteins were first filtered by requiring a P Value<0.05 in a pairwise comparison between the caspase-8 C3FLAG and tagless caspase-8 negative control in either the untreated or TSI treated samples. Known constituents of the native TNFR1 complex 2 (RIPK1, RIPK3, FADD, TRADD and A20) are labelled and highlighted in green while TNKS1 (TNKS) is highlighted in red. P Values calculated using Limma (n = 5). **b,** TNF-induced complex 2 immunoprecipitation using anti-FLAG M2 affinity beads. Western blot analysis of complex 2 and lysates from *Casp8^+/+^, Casp8^+/N3FLAG^* and *Casp8^+/C3FLAG^* BMDMs using the indicated antibodies is shown. Cells were treated with TNF (100 ng/mL) + Smac-mimetic compound A (500 nM) + caspase inhibitor IDN-6556 (5 μM)(TSI) for 1.5 hours before lysis and anti-FLAG immunoprecipitation. FLAG spiked controls contained 3xFLAG peptides at a final concentration of 50 μg/mL. Caspase inhibitor was used to stabilize complex 2. **c-d,** TNF-induced complex 2 immunoprecipitation. Wild-type (WT) BMDMs were treated with TSI (as in **b**) to induce complex 2 assembly. Lysates were immunoprecipitated with anti-RIPK1 (**c**) or anti-cleaved caspase-8 or anti-tankyrase (**d**), separated on SDS/PAGE gels and probed with the indicated antibodies. Filled arrowheads alone denote bands between 100 kDa and 150 kDa detected by anti-tankyrase which might indicate TNKS1 isoform 2 (106 kDa) or TNKS2 (127 kDa). For detailed domain information, see Extended Data Fig. 1d. *indicate IgG chains. Blots are representative of two to three independent experiments.

To further validate these results, we immunoprecipitated endogenous RIPK1 (**Fig. 1c, Extended Data Fig. 1i-j**), FADD (**Extended Data Fig. 1j**) and cleaved caspase-8 (**Fig. 1d, Extended Data Fig. 1i**) from wild-type (WT) BMDMs, MDFs and Mouse Embryonic Fibroblasts (MEFs) and likewise observed TNKS1 co-precipitating with these proteins only when the cells were treated with TSI. Finally, endogenous tankyrases immunoprecipitated FADD, RIPK1 and cleaved caspase-8 from WT BMDMs and MEFs treated with TSI (**Fig. 1d**, **Extended Data Fig. 1i**). TNKS1 also immunoprecipitated with RIPK1, caspase-8 and FADD following TSI treatment of human HT1080 and HT29 cells (**Extended Data Fig. 1k-l**).

Inhibition of protein translation with cycloheximide (CHX) sensitises cells to TNF. TNF+CHX-induced cell death, in contrast to TS-induced death, does not require RIPK1 ^3,19,43–45^. Interestingly, we did not observe the recruitment of TNKS1 to complex 2 after TNF+CHX treatment (**Extended Data Fig. 1m**), suggesting that the different types of cell death are caused by different types of complex 2. Moreover, unlike cIAPs and RIPK1, we did not detect TNKS1 in complex 1 (**Extended Data Fig. 1n**), implying that TNKS1 is specifically recruited together with caspase-8 or FADD to complex 2.

## Complex 2 is PARylated

Tankyrases catalyse the formation of poly-ADP ribose (PAR) chains on their substrates ^41,46^. To determine whether complex 2 becomes PARylated we treated WT BMDMs with TSI ± the tankyrase inhibitor, IWR-1 ^47,48^ and immunoprecipitated PAR chains with an anti-PAR antibody (**Fig. 2a**). Modified RIPK1, indicative of ongoing TNF signalling, precipitated with anti-PAR and was slightly reduced in the presence of IWR-1. Intriguingly, however, cleaved caspase-8 was only immunoprecipitated by the anti-PAR antibody in the absence of the tankyrase inhibitor (**Fig. 2a**).

**Fig. 2.**
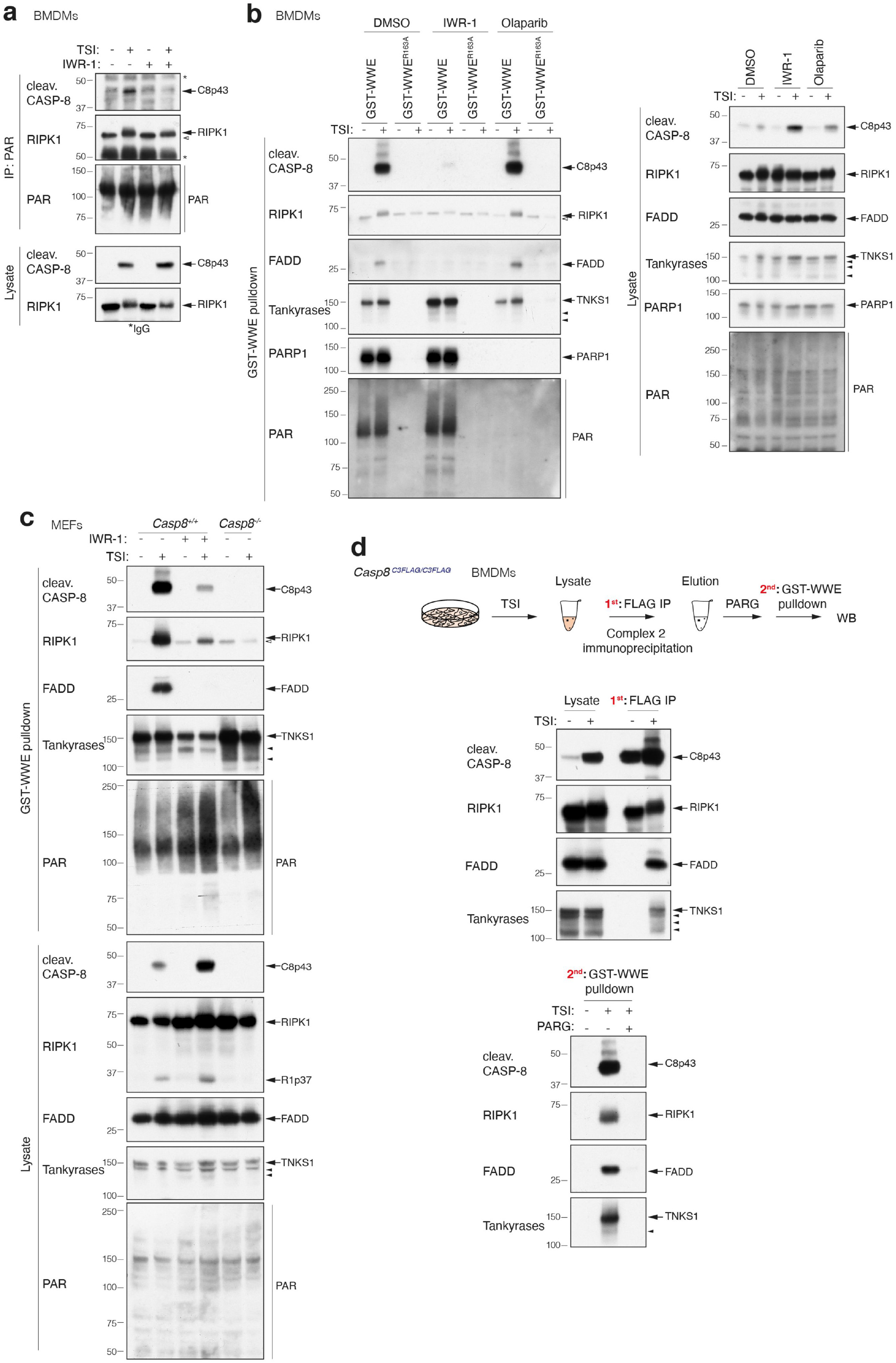
Complex 2 is PARylated. **a,** TNF-induced complex 2 immunoprecipitation using anti-PAR (Trevigen 4335-MC-100). Western blot analysis of complex 2 and lysates from WT BMDMs using the indicated antibodies is shown. Cells were treated with TSI as in Fig. 1 ± tankyrase inhibitor IWR-1 (5 μM) for 1.5 hours before being subjected to anti-PAR immunoprecipitation. **b,** GST-WWE and GST-WWE^R163A^ pulldown of TNF-induced complex 2 from WT BMDMs lysates. Cells were treated with TNF (10 ng/mL) + Smac-mimetic (250 nM) + caspase inhibitor (5 μM) (TSI) ± tankyrase inhibitor IWR-1 (5 μM) or ± PARP1/2 inhibitor olaparib (1μM) for 1.5 hours. Western blot analysis of complex 2 and lysates using the indicated antibodies is shown. **c,** GST-WWE pulldown of TNF-induced complex 2 from *Casp8^+/+^* or *Casp8^+/+^* MEFs lysates. Cells were treated with TSI (as in **a**) ± IWR-1 (10 μM). Western blot analysis of complex 2 and lysates using the indicated antibodies is shown. **d,** Enrichment of PARylated complex 2 using GST-WWE in a sequential pulldown analysis. *Casp8^C3FLAG/C3FLAG^* BMDMs were treated with TSI (as in **a**) and complex 2 was immunoprecipitated using anti-FLAG M2 affinity beads. Immunoprecipitants were eluted with 3x FLAG peptides followed by ± PARG treatment at 37°C for 3 hours before being subjected to GST-WWE pulldown. Western blot analysis of lysates and sequential pulldown using the indicated antibodies is shown. Filled arrowheads alone indicate potential tankyrase species. Empty arrowheads alone denote unmodified RIPK1 that is purified non-specifically by either Sepharose anti-PAR (**a**) or Sepharose GST-WWE (**b-c**). *indicate IgG chains. Blots are representative of two to three independent experiments.

The WWE domain of the E3 ligase RNF146 recognizes the iso-ADP-ribose linkage between two ADP-ribose monomers in PAR chains^49^. We therefore generated a GST fusion of wild-type WWE or a single point mutant (R163A) that is unable to bind PAR chains ^50^ and precipitated lysates from WT BMDMs treated ± TSI ± IWR-1 or the PARP1/2 inhibitor olaparib ^51^ (**Fig. 2b**). Consistent with the anti-PAR immunoprecipitation result, GST-WWE precipitated FADD, modified RIPK1 and cleaved caspase-8 only from lysates of cells treated with TSI (**Fig. 2b**). Notably unmodified RIPK1 was purified using either the wild-type or the mutant WWE motif, suggesting that this interaction and that observed with anti-PAR (**Fig. 2a**) are non-specific and most likely due to the sepharose-beads. As expected TNKS1, PARP1 and PAR chains themselves were all precipitated by GST-WWE but not the GST-WWE R163A mutant in the presence or absence of TSI treatment (**Fig. 2b**). Olaparib treatment substantially reduced the amount of PARP1 and PAR chains precipitated from the lysate, however, consistent with the fact that, at the dose used here it does not inhibit tankyrases, it did not affect the ability of GST-WWE to purify FADD, modified RIPK1 and cleaved caspase-8 (**Fig. 2b**). Conversely, IWR-1 treatment had little impact on the amount of PARP1 and PAR chains precipitated with GST-WWE but almost completely prevented precipitation of FADD, modified RIPK1 and cleaved caspase-8 (**Fig. 2b**). Notably, IWR-1 treatment increased the levels of cleaved caspase-8 in the TSI cell lysates when compared with DMSO control while simultaneously reducing the level of cleaved caspase-8 precipitated by GST-WWE, an effect that was also observed in MEFs (**Fig. 2c**) and MDFs (**Extended Data Fig. 2a**). The precipitation of FADD and modified RIPK1 by GST-WWE upon TSI treatment was completely abrogated by loss of *Casp8* (**Fig. 2c**). To exclude a potential off-target effect of the tankyrase inhibitor IWR-1, and to rule out the possibility that TNKS1 and TNKS2 may compensate for each other ^52^, we depleted TNKS1 in MDFs derived from *Tnks2^-/-^* mice using a doxycycline (Dox)-induced TNKS1 short hairpin RNA (shRNA) and found that the combined absence of TNKS1 and TNKS2 significantly decreased the level of cleaved caspase-8 and modified RIPK1 pulled down with GST-WWE (**Extended Data Fig. 2b**). In contrast, overexpression of TNKS1 isoform 2 but not TNKS2, markedly increased the level of complex 2 components precipitated by GST-WWE (**Extended Data Fig. 2c**), indicating that TNKS1 plays a predominant role in complex 2 PARylation.

To confirm that at least one complex 2 component was PARylated we performed a FLAG immunoprecipitation from homozygous *Casp8 ^C3FLAG/C3FLAG^* BMDMs stimulated with TSI and then treated this ± poly-ADP ribose glycohydrolase (PARG), a dePARylating enzyme that cleaves conjugated ADP-ribose polymers ^53,54^ (**Fig. 2d**). GST-WWE was then used to sequentially purify PARylated proteins from the purified complex. Consistent with our previous results, FADD, modified RIPK1, TNKS1 and cleaved caspase-8 were precipitated following TSI treatment but only if PAR chains had not been removed by PARG treatment (**Fig. 2d**). A similar approach using the ADP-ribose binding macrodomain *Af1521* from *Archaeoglobus fulgidus,* which binds mono-ADP-ribose groups and the terminal ribose in PAR chains ^49^, also sequentially precipitated FADD, modified RIPK1, TNKS1 and cleaved caspase-8 (**Extended Data Fig. 2d**).

WWE domains are found in many E3 ubiquitin ligases ^55^, including HUWE1 and TRIP12. The critical residues for PAR binding are conserved in most WWE domains and HUWE1 and TRIP12 WWE domains specifically interact with PAR chains ^56^. To determine whether there might be some specificity to the complex 2 interaction, we performed a PAR pulldown assay using GST-HUWE1, -TRIP12 and -RNF146 WWE fusion proteins (**Extended Data Fig. 2e**). GST-RNF146 WWE was more efficient than GST-HUWE1 WWE which in turn was far more efficient than GST-TRIP12, at precipitating complex 2 components, suggesting that there may be some specificity and indicating that the RNF146 WWE is optimal for PARylated complex 2 purification (**Extended Data Fig. 2e**).

## Tankyrases limit TNF-induced cell death

Thus far, our data suggested that TNKS1 is a functional component of complex 2 and complex 2 undergoes PARylation and also hinted that ADP-ribosylation might limit caspase-8 activation. To explore this further we treated WT BMDMs with increasing doses IWR-1 and measured TNF-induced cell death by flow cytometry. Consistent with our earlier Western blot analyses (**Fig. 2**), BMDMs were rendered increasingly sensitive to TNF plus Smac-mimetic-induced apoptosis (TS) ^1,15,30,31,38,57,58^ and TSI-induced necroptosis ^34,59–70^ by increasing doses of IWR-1 (**Fig. 3a-b**). This sensitisation was reversed by inhibition of RIPK1 kinase activity with necrostatin-1s, suggesting that tankyrase inhibition sensitised cells to TNF-induced cell death in a RIPK1 kinase-dependent manner (**Fig. 3a-b**). Inhibition or depletion of tankyrases also sensitized MDFs to TS-induced death (**Extended Data Fig. 3a-b**), but consistent with the lack of TNKS1 in TNF+CHX-induced complex 2 (**Extended Data Fig. 1m**), inhibition of tankyrases did not affect TNF+CHX-induced cell death (**Extended Data Fig. 3a**). Another tankyrase inhibitor, Az6102 ^51^, also increased sensitivity to TNF-induced death, while the PARP1/2 inhibitor, olaparib, did not (**Extended Data Fig. 3c**). Consistent with the increased cell death, increasing IWR-1 concentrations increased the levels of cleaved caspase-8 and caspase-3 (**Fig. 3c**) observed in TS treated BMDMs and phospho-RIPK3 and phospho-MLKL in TSI treated cells (**Fig. 3d**). The clinical Smac-mimetic birinapant kills leukemic cells in a TNF-dependent manner^30,31,38^, and consistent with this, and our previous data, MLL-AF9/NRas^G12D^ cells were dramatically sensitised to both apoptotic and necroptotic cell death by increasing doses of IWR-1 (**Extended Data Fig. 3d)**.

**Fig. 3.**
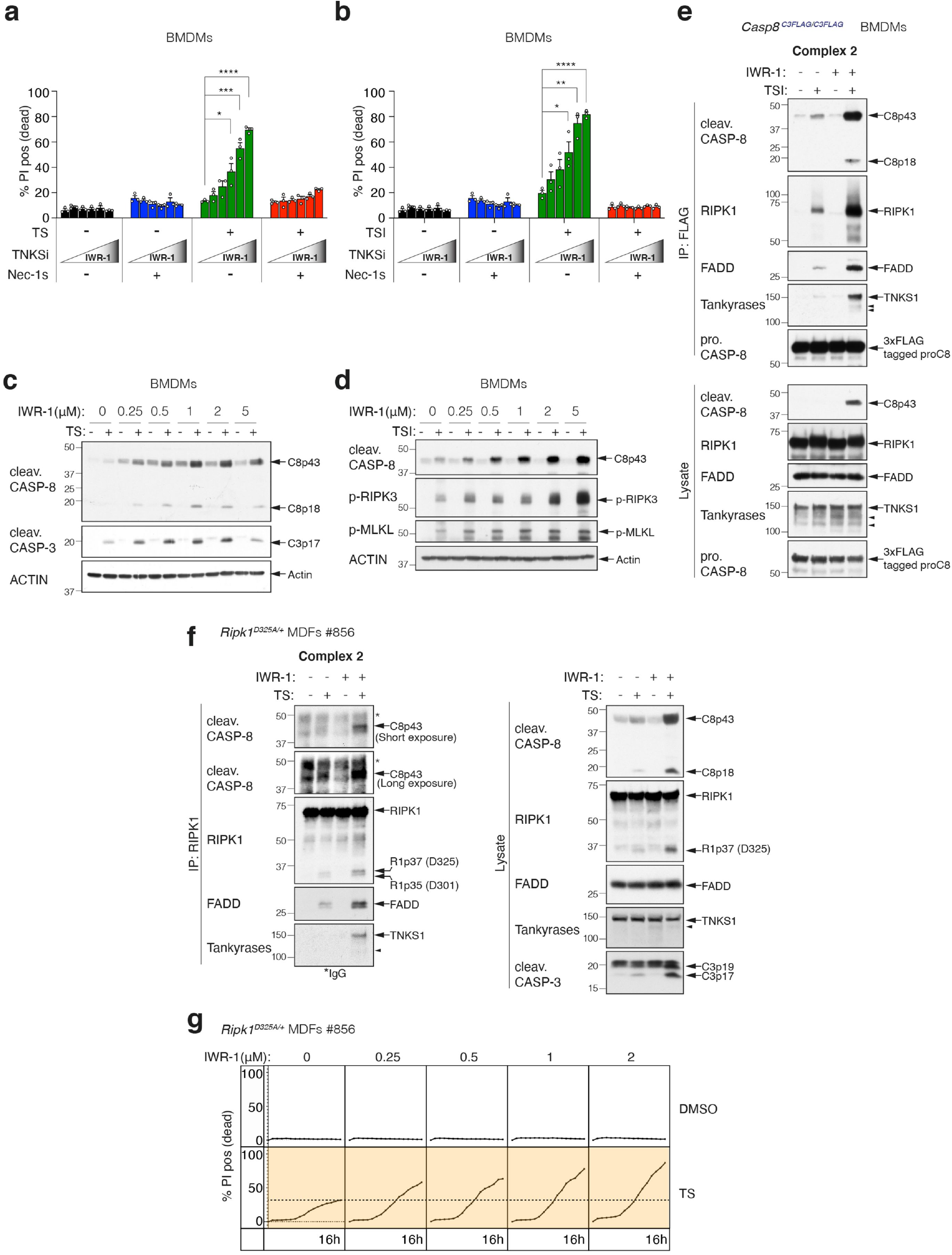
Tankyrases limit TNF-induced cell death. **a,** Level of cell death assessed by propidium iodide (PI) positive cells. WT BMDMs were treated with TNF (10 ng/mL) + Smac-mimetic (500 nM) (TS) ± IWR-1 (250nM, 500nM, 1 μM, 2 μM, 5 μM) ± Nec-1s (10 μM) for 24 hours. Graphs show mean ± SEM, n=3 biologically independent repeats. Comparisons were performed with a Student’s t test whose values are denoted as *p ≤ 0.05, ***p ≤ 0.001 and ****p ≤ 0.0001. **b,** Level of cell death assessed by PI positive cells. WT BMDMs were treated with TNF (10 ng/mL) + Smac-mimetic (10 nM) + caspase inhibitor (5 μM) (TSI) ± IWR-1 (250nM, 500nM, 1 μM, 2 μM, 5 μM) ± Nec-1s (10 μM) for 16 hours. Graphs show mean ± SEM, n=3 biologically independent repeats. Comparisons were performed with a Student’s t test whose values are denoted as *p ≤ 0.05, **p ≤ 0.01 and ****p ≤ 0.0001. **c,** Western blot analysis of cell lysates from WT BMDMs using indicated antibodies is shown. Cells were treated with TNF (10 ng/mL) + Smac-mimetic (500 nM) (TS) ± IWR-1 (250nM, 500nM, 1 μM, 2 μM, 5 μM) for 8 hours. **d,** Western blot analysis of cell lysates from WT BMDMs using indicated antibodies is shown. Cells were treated with TNF (10 ng/mL) + Smac-mimetic (20 nM) + caspase inhibitor (5 μM) (TSI) ± IWR-1 (250nM, 500nM, 1 μM, 2 μM, 5 μM) for 8 hours. **e,** TNF-induced complex 2 immunoprecipitation using anti-FLAG M2 affinity beads. Western blot analysis of complex 2 and lysates from *Casp8*^C3FLAG/C3FLAG^ BMDMs using the indicated antibodies is shown. Cells were treated with TNF (10 ng/mL) + Smac-mimetic (50 nM) + caspase inhibitor (5 μM) (TSI) ± IWR-1 (5 μM) for 1.5 hours before being subjected to anti-FLAG immunoprecipitation. **f,** TNF-induced complex 2 immunoprecipitation using anti-RIPK1 antibody. Western blot analysis of complex 2 and lysates from *Ripk1^D325A/+^* heterozygous MDFs using the indicated antibodies is shown. Cells were treated with TNF (50 ng/mL) + Smac-mimetic (100 nM) ± IWR-1 (5 μM) for 2 hours before being subjected to anti-RIPK1 immunoprecipitation. **g,** Cell death monitored by time-lapse imaging of PI staining over 16 hours using IncuCyte. *Ripk1^D325A/+^* heterozygote MDFs were treated with TNF (50 ng/mL) + Smac-mimetic (25 nM) (TS) ± IWR-1 (250nM, 500nM, 1 μM, 2 μM) for 16 hours. Dashed lines denote the PI count without IWR-1 treatment for reference. Results from two additional, biologically independent MDFs are shown in Extended Data Fig. 3f. Filled arrowheads alone indicate potential tankyrase species. *indicate IgG chains. Blots are representative of two to three independent experiments.

To determine why cells were more sensitive to TNF-induced cell death when tankyrase activity was inhibited we immunoprecipitated complex 2 from *Casp8^C3FLAG/C3FLAG^* BMDMs and MEFs treated with TSI ± IWR-1. By selecting a TSI dose that induced only low levels of caspase-8 activation, we were able to show that tankyrase inhibition dramatically increased the amount of complex 2 that could be immunoprecipitated by anti-FLAG beads, suggesting that tankyrase-mediated ADP-ribosylation reduces the stability of complex 2 (**Fig. 3e, Extended Data Fig. 3e**). Typically, complex 2 is difficult to purify unless a caspase inhibitor, such as emricasan/IDN-6556, is used to stabilise it ^3,38^. However, this makes it difficult to test whether tankyrase inhibition increases complex 2 formation in the absence of a caspase inhibitor. To circumvent this issue, we took advantage of the fact that complex 2 can be isolated more readily from cells expressing an uncleavable form of RIPK1 ^71^. We therefore treated *Ripk1^D325A/+^* heterozygous MDFs with TS ± IWR-1 and immunoprecipitated RIPK1 and found that tankyrase inhibition also increased the amount of complex 2 that could be purified from these cells and sensitised them to TNF-induced cell death in a dose dependent manner (**Fig. 3f-g, Extended Data Fig. 3f**).

## The tankyrase-RNF146 axis regulates the stability of complex 2 and TNF-induced death

Tankyrases regulate a number of other signalling pathways^48,72–75^, and the most well-studied is the Wnt pathway where tankyrase-mediated ADP-ribosylation of Axin recruits the E3 ligase RNF146 via its WWE motif. RNF146 then ubiquitylates Axin causing its recruitment to and degradation by the proteasome ^50,56,76–78^. Given the increased stability of complex 2 in the presence of tankyrase inhibitor IWR-1 that we observed, we hypothesized that tankyrase-mediated ADP-ribosylation of complex 2 might function analogously to recruit RNF146 and promote its proteasomal degradation. In accord with this hypothesis RNF146 was recruited to complex 2 immunoprecipitated from *Casp8^+/C3FLAG^* heterozygote MEFs treated with TSI (**Fig. 4a**). Furthermore, there was a reduction in the precipitation of ubiquitylated complex 2 components using a GST-UBA fusion protein, when cells were treated with IWR-1 (**Fig. 4b**). Consistent with the idea that proteasomal mediated degradation limits complex 2 levels, we observed a striking increase in the amount of ubiquitylated complex 2 when cells were treated with the proteasomal inhibitor MG132 (**Fig. 4c**). To avoid the possibility that constitutive loss of RNF146 affected cell viability, we generated stable Dox inducible RNF146 shRNA expressing cells and immunoprecipitated RIPK1 in the presence or absence of Dox. Similarly to the proteasome inhibitor experiment, we saw that there was a stark increase in the levels of complex 2 in the cells with reduced levels of RNF146 when compared with control shRNA expressing cells (**Fig. 4d**), and as expected shRNF146 expressing cells were more sensitive to TNF-induced cell death (**Fig. 4e, Extended Data Fig. 4**).

**Fig. 4.**
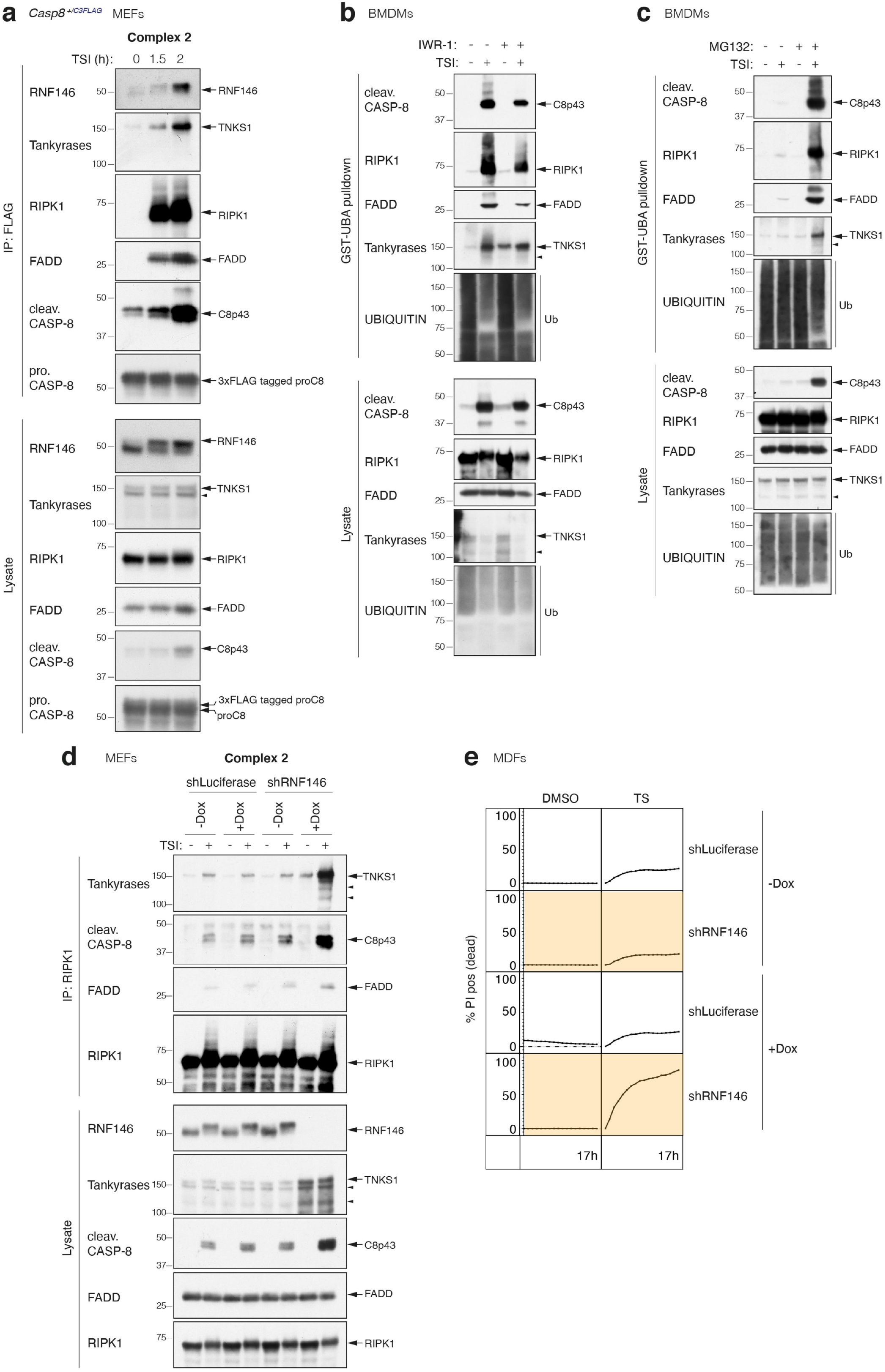
The tankyrase-RNF146 axis regulates the stability of complex 2 and TNF-induced death. **a,** TNF-induced complex 2 immunoprecipitation using anti-FLAG M2 affinity beads. Western blot analysis of complex 2 and lysates from *Casp8^+/C3FLAG^* MEFs using the indicated antibodies is shown. Cells were treated with TNF (100 ng/mL) + Smac-mimetic (500 nM) + caspase inhibitor (5 μM) (TSI) for the indicated timepoints before being subjected to anti-FLAG immunoprecipitation. **b,** GST-UBA pulldown of TNF-induced complex 2 from WT BMDM lysates. Cells were treated with TNF (100 ng/mL) + Smac-mimetic (500 nM) + caspase inhibitor (5 μM) (TSI) for 1.5 hours ± IWR-1 (5 μM). Western blot analysis of complex 2 and lysates using the indicated antibodies is shown. **c,** GST-UBA pulldown of TNF-induced complex 2 from WT BMDMs lysates. Cells were pre-treated with ± proteasome inhibitor MG132 (10 μM) for 2 hours, followed by TNF (10 ng/mL) + Smac-mimetic (50 nM) + caspase inhibitor (5 μM) (TSI) ± MG132 (10 μM) for another 2 hours. Western blot analysis of complex 2 and lysates using the indicated antibodies is shown. **d,** TNF-induced complex 2 immunoprecipitation using anti-RIPK1 antibody. WT MEFs expressing Dox-inducible shLuciferase or shRNF146 were pre-treated with ± Dox (1μg/mL) for 48 hours. Cells were then treated with TNF (100 ng/mL) + Smac-mimetic (500 nM) + caspase inhibitor (5 μM) (TSI) ± Dox (1 μg/mL) for another 2 hours. Western blot analysis of complex 2 and lysates using the indicated antibodies is shown. **e,** Cell death monitored by time-lapse imaging of PI staining over 17 hours. WT MDFs expressing Dox-inducible shLuciferase or shRNF146 were pre-treated with ± Dox (1μg/mL) for 48 hours, followed by TNF (100 ng/mL) + Smac-mimetic (50 nM) (TS) ± Dox (1μg/mL) for another 17 hours. n=1 biological repeat. Filled arrowheads alone indicate potential tankyrase species. Blots are representative of two to three independent experiments.

## Viral macrodomains sensitise TNF-induced death

TNF is an important part of the mammalian anti-pathogen armamentarium and as a consequence is frequently targeted by pathogens which produce proteins that interfere with the pathway^2^. The TNF pathway has however several mechanisms to respond to interference and one of those is to trigger cell death. This begs the question whether ADP-ribosylation of complex 2 also serves to control for interference and whether the increased death that we observed when tankyrase activity is inhibited might mimic some form of pathogen manipulation. A number of viruses, including Coronaviruses, express evolutionarily conserved MacroD type macrodomains ^79,80^, similar to that of *Af1521* that we used to precipitate complex 2, that are able to bind to mono-ADP-ribosylated proteins or to the end of poly-ADP-Ribose chains and in some cases have been shown to remove ADP-ribose from mono-ADP-ribosylated proteins ^81–85^. We therefore asked whether inducible expression of the macrodomain from SARS-CoV-2 or a closely related VEEV macrodomain might affect TNF-induced cell death. Consistent with the idea that ADP-ribosylation of complex 2 could serve as a checkpoint to detect perturbations in TNF signalling we found that expression of both these viral macrodomains markedly increased the sensitivity of cells to TNF-induced cell death (**Fig. 5**).

**Fig. 5.**
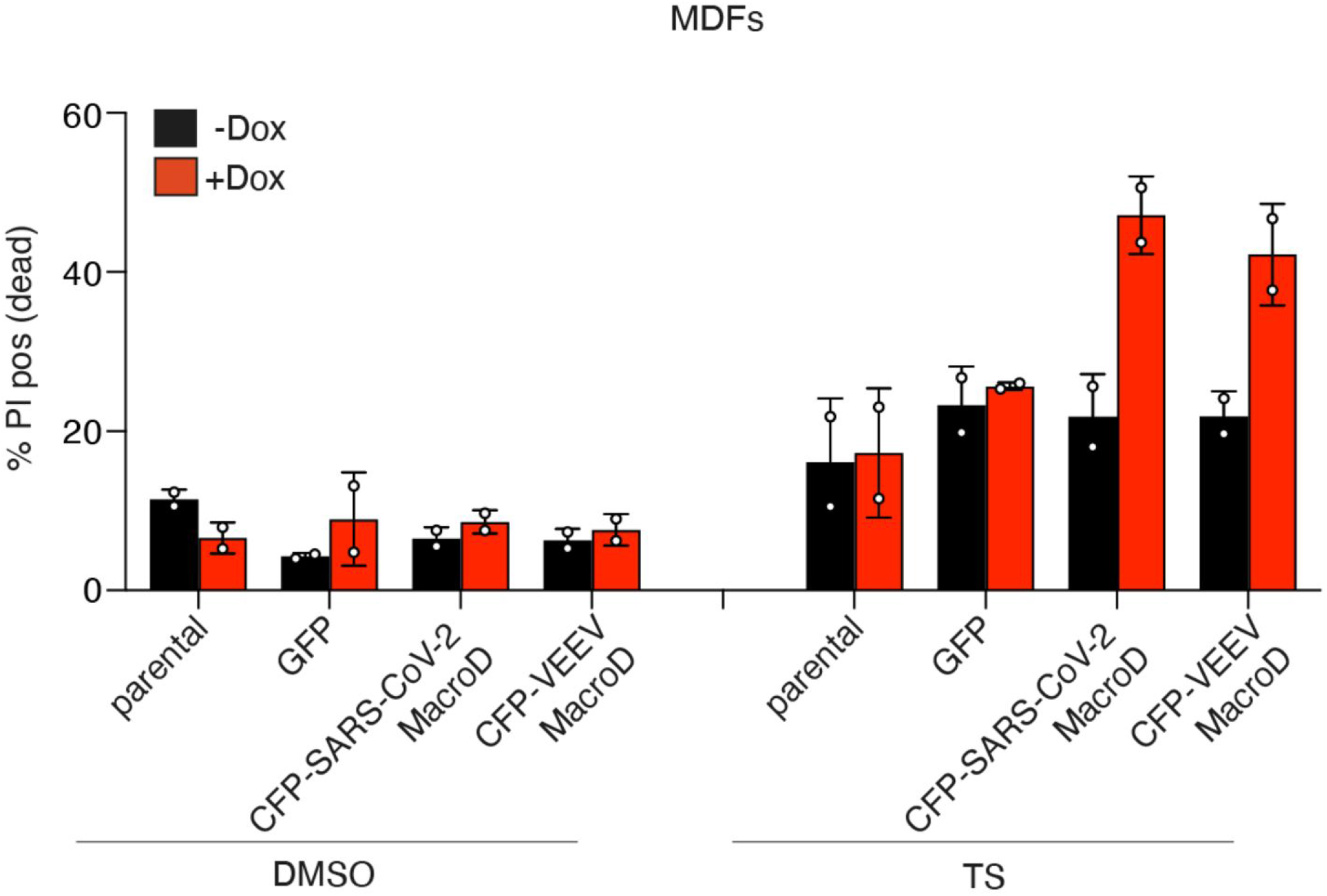
Viral macrodomains sensitise cells to TNF-induced death. Parental WT MDFs and MDFs expressing Dox-inducible GFP control or CFP-SARS-CoV-2 macrodomain or CFP-VEEV macrodomain were pre-treated with ± Dox (10ng/mL) for 3 hours. Cells were then treated with TNF (50 ng/mL) + Smac-mimetic (10 nM) (TS) in the absence of Dox for another 20 hours, and amount of cell death was assessed by PI staining and flow cytometry. Graphs show mean ± SD throughout, n = 2 independent biological repeats.

We show that the ability of TNF to induce cell death is regulated by tankyrase-mediated PARylation. Interestingly, while TNKS1 was readily recruited to complex 2 upon Smac-mimetic treatment, it was not detectable in complex 2 assembled in response to cycloheximide. This suggests that ADP-ribosylation is a context sensitive regulator and since RIPK1 involvement is a major difference in these two complexes, it suggests RIPK1 might be directly involved.

Tankyrase 1 & 2 regulate a number of signalling pathways and one possibility is that the PARylation-mediated by tankyrases might allow different signalling pathways to interact and co-ordinate with one another. In particular there is evidence linking TNF signalling with the Wnt and GSK3 signalling pathways as well as cell cycle and cell division, all of which are known to be regulated by tankyrases ^46,86,87^. Indeed, specific and potent tankyrase inhibitors, such as IWR-1, were developed to block Wnt signalling in cancers yet clearly sensitise cells to TNF killing and this unintended activity might increase the efficacy of these drugs in tumors with an inflammatory component. Furthermore, it has been noted that some cancers are sensitive to these inhibitors without apparently affecting Wnt signalling thus opening up the possibility that sensitivity to TNF might be an additional predictive biomarker to consider when using these drugs.

Despite its defensive intent, excessive TNF-induced cell death can cause serious pathology, and SARS-CoV-2 infection triggers caspase-8 activation and apoptosis in mice and the postmortem lung sections of COVID-19 patients also contain markers of extrinsic TNF-induced apoptosis ^88,89^. Viral macrodomains have been shown to either bind to or hydrolyse ADP-ribose^81,85^, and since inducible expression of the macrodomains of SARS-CoV-2 and VEEV sensitised cells to TNF-induced cell death, this suggests that ADP-ribosylation may serve as yet another mechanism to allow TNF to retaliate against a dangerous infection by inducing cell death. This idea is supported by the observation that PARP-10, a mono-ADP-ribosyltransferase, inhibits IL-1β/TNF-induced NF-κB signalling^90^.Given the broad involvement of ADP-ribosylation in other signalling pathways ^91,92^, one intriguing possibility is that pathogens select for the ability to interfere with ADP-ribosylation to target these pathways and that ADP-ribosylation has been co-opted into the TNF response to control for the integrity of these pathways rather than of the TNF pathway alone.

## EXTENDED DATA FIGURE LEGENDS

**Extended Data Fig. 1.**
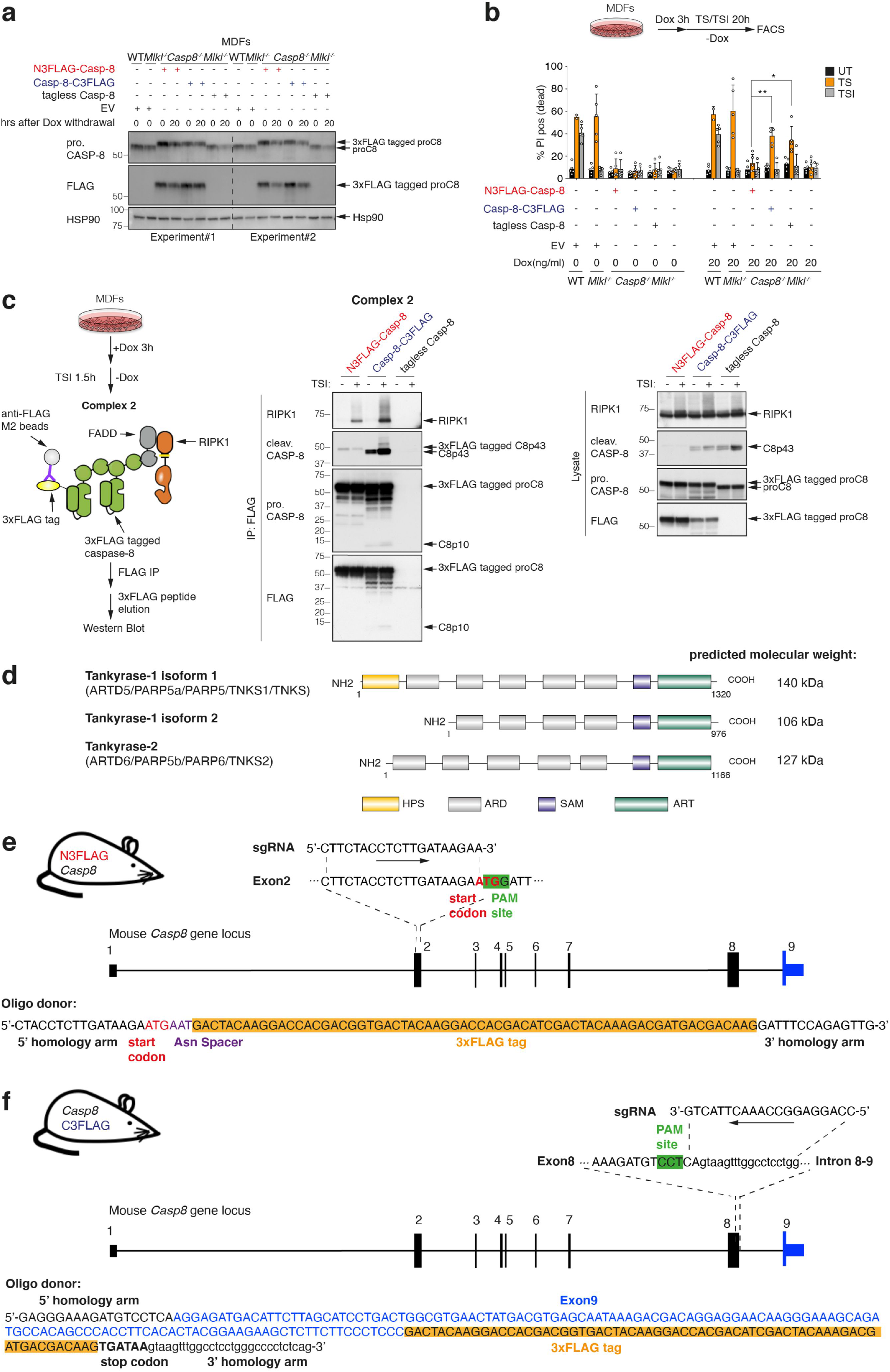
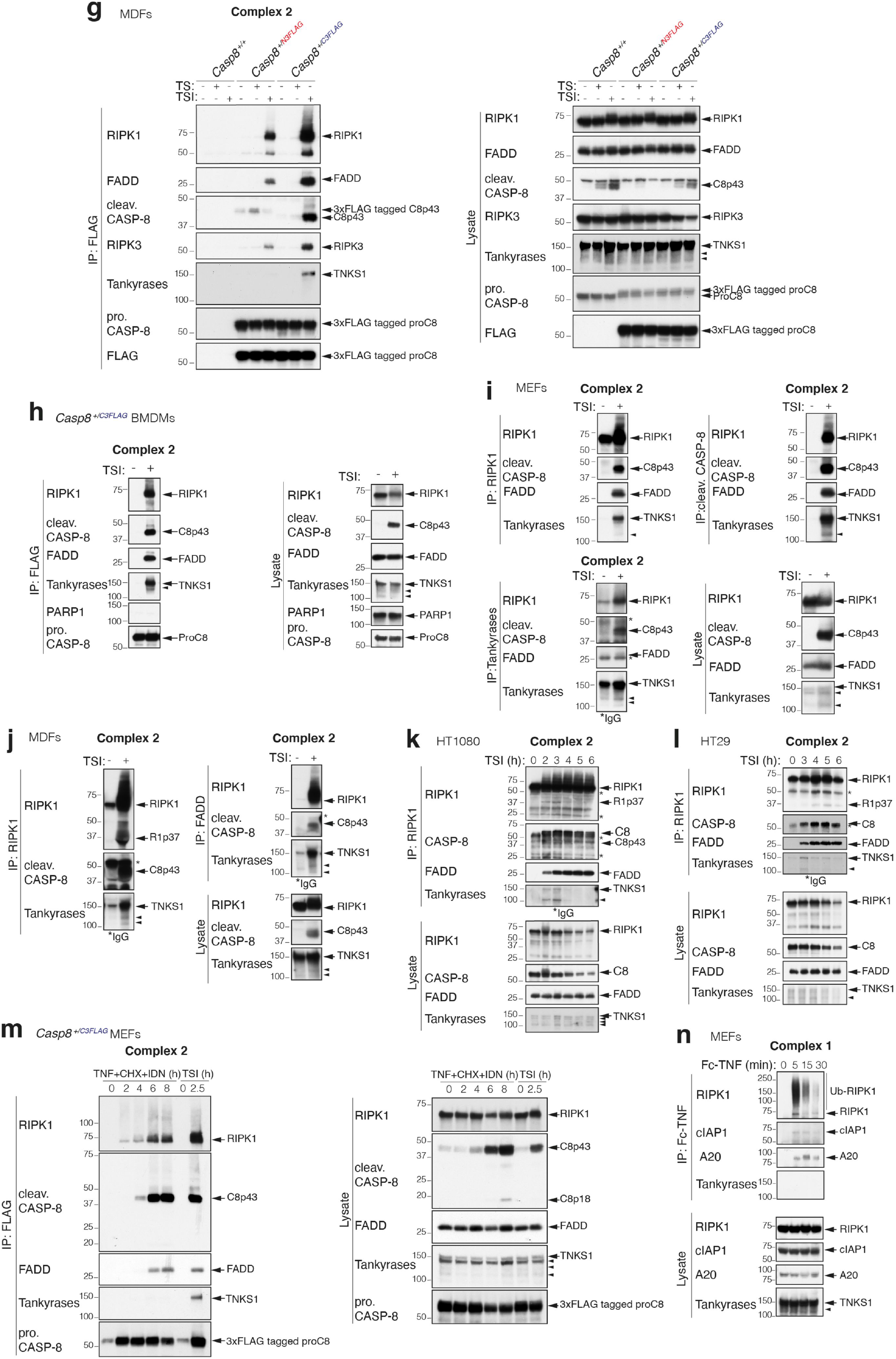
Tankyrase-1 is a novel interactor of native TNFR1 complex 2. **a,** Western blot analysis of cell lysates from *Casp8^-/-^.Mlkl^-/-^* MDFs expressing doxycycline (Dox)-inducible N- (red) or C- (blue) 3x FLAG tagged murine caspase-8 or tagless caspase-8. Wild-type (WT) or *Mlkl^-/-^* MDFs expressing an empty vector (EV) were used as controls. Cells were treated with 20 ng/mL Dox for 3 hours and then Dox was withdrawn. Samples were harvested 0 hour or 20 hours after Dox withdrawal for Western blot analysis. **b,** Level of cell death assessed by propidium iodide (PI) positive cells. Cells were pre-treated with 20 ng/mL Dox for 3 hours followed by stimulation with TNF (100 ng/mL) + Smac-mimetic compound A (500 nM) (TS) ± caspase inhibitor IDN-6556 (5 μM) for 20 hours in the absence of Dox. Graphs show mean ± SD, n=3 independent experiments. Comparisons were performed with a Student’s t test whose values are denoted in the figures as *p ≤ 0.05 and **p ≤ 0.01. **c,** Left, schematic depicting the anti-FLAG immunoprecipitation. Right, TNF-induced complex 2 immunoprecipitation using anti-FLAG M2 affinity beads. Western blot analysis of complex 2 from *Casp8^-/-^.Mlkl^-/-^* MDFs expressing Dox-inducible N- or C-3x FLAG tagged murine caspase-8 or tagless caspase-8 using the indicated antibodies. Cells were treated with 20 ng/mL Dox for 3 hours followed by stimulation with TNF (100 ng/mL) + Smac-mimetic (500 nM) + caspase inhibitor (5 μM) (TSI) for 1.5 hours in the absence of Dox before subjected to anti-FLAG immunoprecipitation. Caspase inhibitor was used to stabilize complex 2. The Western blot is representative of five independent experiments. **d,** Schematic comparison of the domain architecture of the murine TNKS1, TNKS1 isoform2 and TNKS2. Domains are: HPS: histidine, proline and serine-rich region; ARD: ankyrin repeat domains; SAM: sterile α-motif; ART: poly(ADP-ribose) polymerases catalytic domain. ARDs provide binding sites for interaction between tankyrases and other proteins. The SAM domain mediates protein-protein interactions, form homo- and hetero-oligomers and also binds to DNA, RNA and lipids. SAM domain is also critical for optimal catalytic activity. The ART domain is responsible for the ADP-ribosyltransferase activity. **e-f,** Schematic representation of the generation of *Casp8*^N3FLAG^ (**e**) or *Casp8*^C3FLAG^ (**f**) mice using CRIPSR/Cas9 technology. For N-3x FLAG tagged caspase-8 knock-in mice, an Asn Spacer was introduced into the oligo donor to ensure successful gene translation. For C-3x FLAG tagged caspase-8 knock-in mice, the PAM site was in exon 8 and an oligo donor composed of protein coding region of exon 9 with a 3x FLAG tag followed by two stop codons were designed because there was no usable PAM site at the last exon (exon 9) of *Casp8* gene and a *Casp8* pseudogene known as Gm20257 showed ~133 bp of sequence identity to *Casp8* exon 9 and was nearby on the same chromosome (chromosome 1). **g-h,** TNF-induced complex 2 immunoprecipitation using anti-FLAG M2 affinity beads. Western blot analysis of complex 2 and lysates from *Casp8*^+/+^, *Casp8*^+/N3FLAG^ and *Casp8*^+/C3FLAG^ MDFs **(g)** or *Casp8*^+/C3FLAG^ BMDMs **(h)** using the indicated antibodies is shown. Cells were treated with TNF (100 ng/mL) + Smac-mimetic (500 nM) with or without caspase inhibitor (5 μM) for 1.5 hours before being subjected to anti-FLAG immunoprecipitation. Caspase inhibitor was used to stabilize complex 2. **i-j,** TNF-induced complex 2 immunoprecipitation. WT MEFs (**i**) or MDFs (**j**) were treated with TSI (as in **g-h**) to induce complex 2 assembly. The lysates were immunoprecipitated with anti-RIPK1 or anti-cleaved caspase-8 or anti-FADD or anti-tankyrase. Western blot analysis using the indicated antibodies is shown. **k-l,** TNF-induced complex 2 immunoprecipitation using anti-RIPK1. Western blot analysis of complex 2 and lysates from HT1080 (**k**) and HT29 (**l**) cells using the indicated antibodies is shown. Cells were treated with TSI (as in **g-h**) for indicated time points. **m,** TNF-induced complex 2 immunoprecipitation using anti-FLAG M2 affinity beads. Western blot analysis of complex 2 and lysates from *Casp8*^+/C3FLAG^ MEFs using the indicated antibodies is shown. Cells were treated with TNF (100 ng/mL) + Smac-mimetic (500 nM) + caspase inhibitor (5 μM) (TSI) for 2.5 hours or TNF (100 ng/mL) + cycloheximide (CHX) (1μg/mL) + caspase inhibitor (5 μM) (TNF+CHX+IDN) for the indicated time points, followed by immunoprecipitation with anti-FLAG M2 affinity beads. **n,** TNF-induced complex 1 immunoprecipitation. WT MEFs were treated with Fc-TNF (1 μg/mL) for the indicated time points, followed by immunoprecipitation with protein A Sepharose and Western blot analysis. Filled arrowheads alone indicate potential tankyrase species. *indicate IgG chains. Blots are representative of two to three independent experiments.

**Extended Data Fig. 2.**
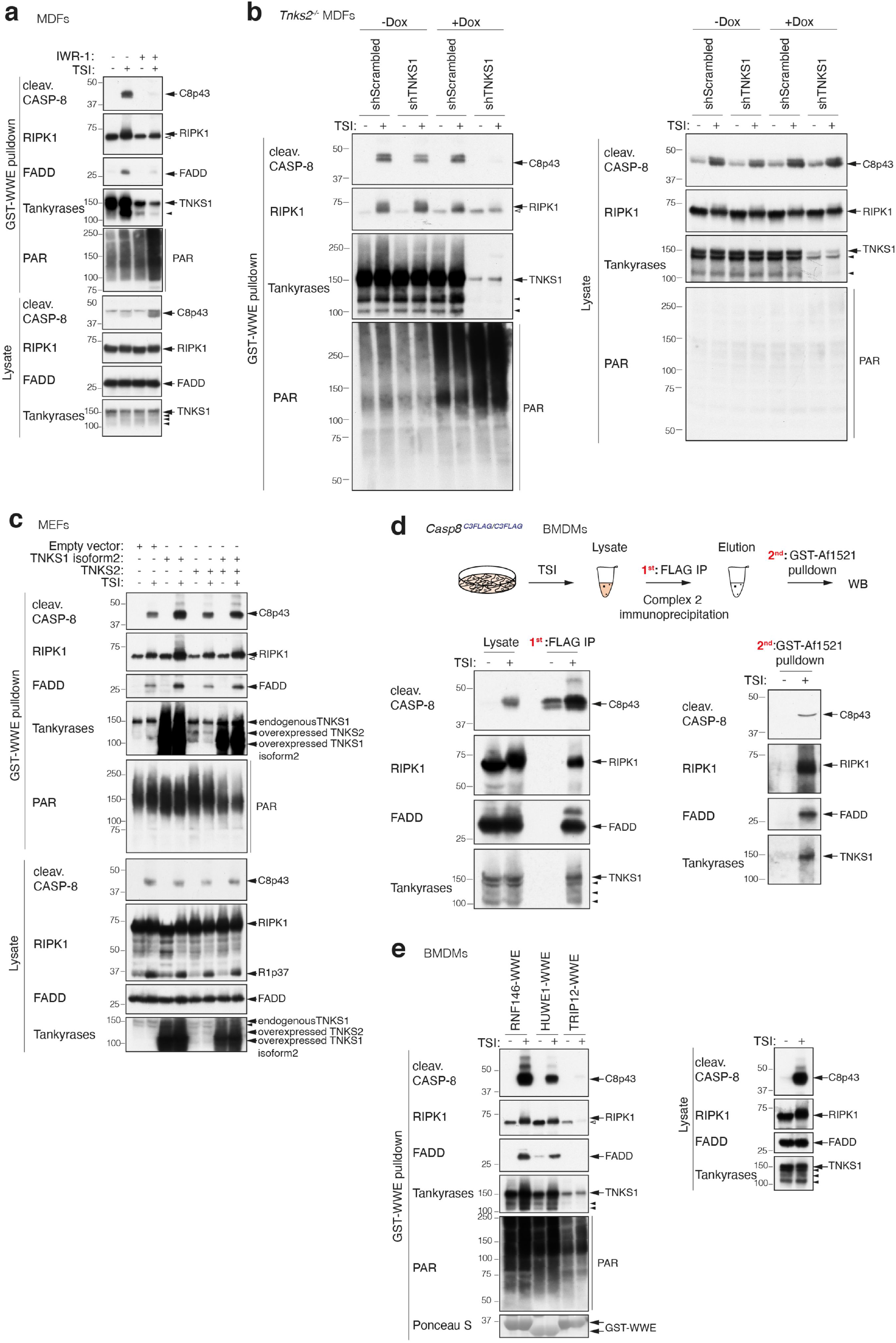
Complex 2 is PARylated. **a,** GST-WWE pulldown of TNF-induced complex 2 from WT MDF lysates. Cells were treated with TNF (100 ng/mL) + Smac-mimetic (500 nM) + caspase inhibitor (5 μM) (TSI) for 1.5 hours ± tankyrase inhibitor IWR-1 (10 μM). Western blot analysis of complex 2 and lysates using the indicated antibodies is shown. **b,** GST-WWE pulldown of TNF-induced complex 2. *Tnks2^-/-^* MDFs expressing Dox-inducible Scrambled shRNA or TNKS1 shRNA were pre-treated with ± Dox (1μg/mL) for 48 hours before being stimulated with TNF (100 ng/mL) + Smac-mimetic (500 nM) + caspase inhibitor (5 μM) (TSI) ± Dox (1μg/mL) for 1.5 hours. Cell lysates were subjected to GST-WWE pulldown. Western blot analysis of complex 2 and lysates using the indicated antibodies is shown. **c,** GST-WWE pulldown of TNF-induced complex 2. WT MEFs expressing Dox-inducible murine TNKS1 (isoform 2) or/and TNKS2 or empty vector were pre-treated with 20 ng/mL Dox overnight before being stimulated with TSI (as in **b**) for 1.5 hours. Cell lysates were subjected to GST-WWE pulldown. Western blot analysis of complex 2 and lysates using the indicated antibodies is shown. **d,** Enrichment of PARylated complex 2 using GST-Af1521 in a sequential pulldown analysis. *Cas*^C3FLAG/C3FLAG^ BMDMs were treated with TSI (as in **b**) and complex 2 was immunoprecipitated using anti-FLAG M2 affinity beads. Immunoprecipitants were eluted using 3x FLAG peptides followed by GST-Af1521 pulldown. Western blot analysis of lysates and sequential pulldown using the indicated antibodies is shown. **e,** GST-HUWE1, -TRIP12 and -RNF146 WWE pulldown of TNF-induced complex 2 from WT BMDMs. Cells were treated with TSI (as in **b**) and lysates were subjected to GST-WWE pulldown assays. Western blot analysis of complex 2 and lysates using the indicated antibodies is shown. Ponceau S staining of the purified proteins and their quantities used in the pulldown assay is shown. Filled arrowheads alone indicate potential tankyrase species. Empty arrowheads alone denote unmodified RIPK1 that is purified non-specifically by Sepharose GST-WWE. Blots are representative of two to three independent experiments.

**Extended Data Fig. 3.**
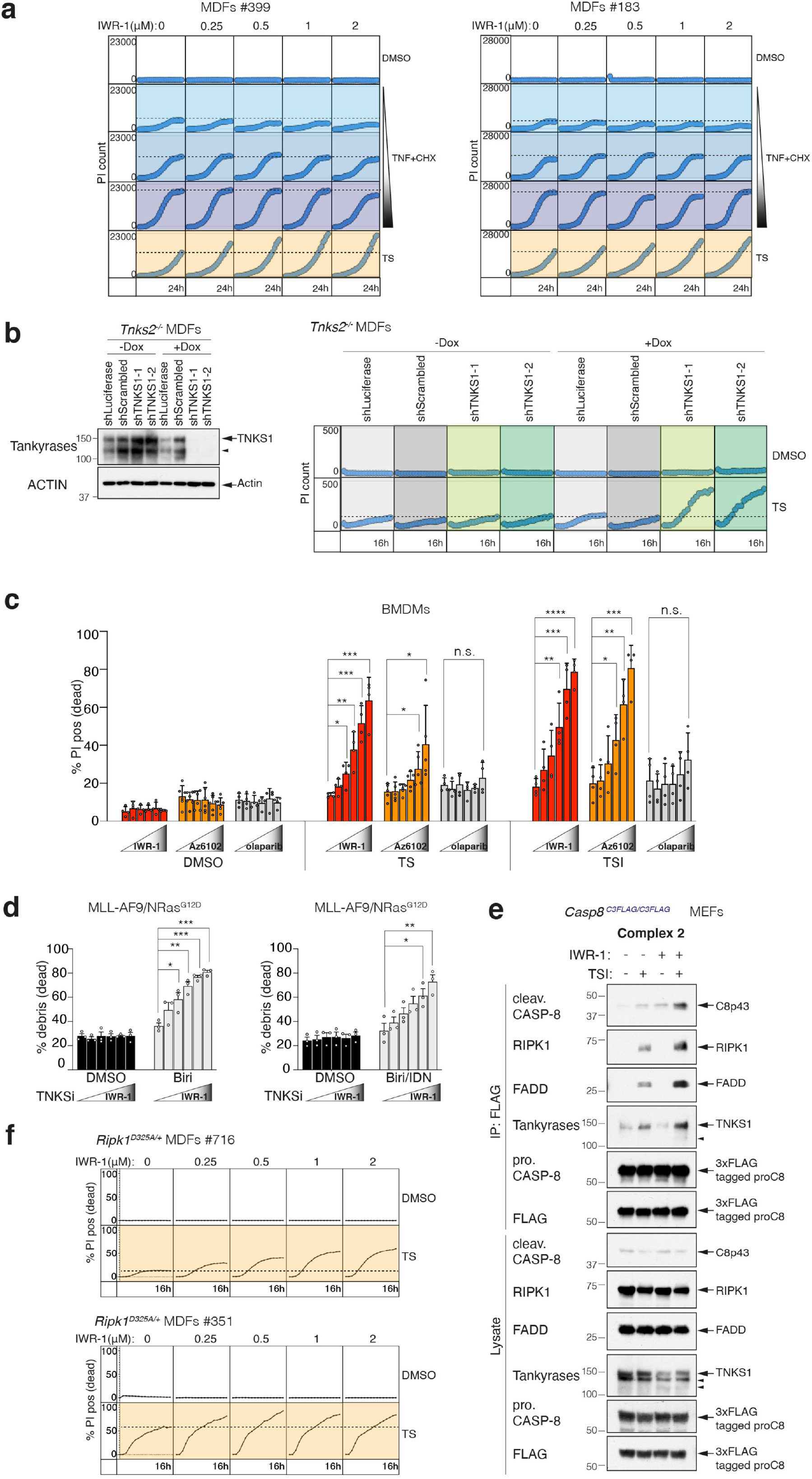
Tankyrases limit TNF-induced cell death. **a,** Cell death of WT MDFs, monitored by time-lapse imaging of PI staining (dead cells) over 24 hours. WT MDFs were treated with DMSO, TNF+cycloheximide (TNF+CHX) or TNF+Smac-mimetic (TS) (rows) ± tankyrase inhibitor IWR-1 (columns) for 24 hours. TNF: 50 ng/mL. Smac-mimetic: 50nM. CHX: 0.25μg/mL, 0.5μg/mL, 1μg/mL. IWR-1: 250nM, 500nM, 1 μM, 2 μM. Cell death was quantified by PI uptake and time-lapse imaging every 1 hour using IncuCyte. Dashed lines denote the PI count without IWR treatment for reference. The results from two biologically independent MDF lines are shown. **b,** Left, Western blot analysis of TNKS1 knockdown efficiency in *Tnks2^-/-^* MDFs expressing Dox-inducible shLuciferase, shScrambled or two independent TNKS1 shRNA. Cells were pre-treated with ± Dox (1μg/mL) for 48 hours and then subjected to Western blot analysis. Filled arrowhead alone indicates potential tankyrase species. Right, *Tnks2*^-/-^ MDFs expressing Dox-inducible shLuciferase, shScrambled or two independent TNKS1 shRNAs were pre-treated with ± Dox (1μg/mL) for 48 hours followed by TNF (10 ng/mL) + Smac-mimetic (25 nM) (TS) ± Dox (1μg/mL) for 16 hours. Cell death was quantified by PI uptake and time-lapse imaging every 1 hour using IncuCyte. Dashed lines denote the PI count in cells where the shRNA is not induced for reference. n=1 biological repeat. **c,** Amount of cell death assessed by PI positive cells by flow cytometry. WT BMDMs were treated with TNF (10 ng/mL) + Smac-mimetic (500 nM) (TS) or TNF (10 ng/mL) + Smac-mimetic (10 nM) + caspase inhibitor (5 μM) (TSI) ± tankyrase inhibitor IWR-1 or ± Az6102 or ± PARP1/2 inhibitor olaparib for 16 hours. IWR-1: 250nM, 500nM, 1 μM, 2 μM, 5 μM. Az6102: 125nM, 250nM, 500nM, 1 μM, 2 μM. Olaparib: 62.5nM, 125nM, 250nM, 500nM, 1 μM. Graphs show mean ± SEM, n = 4-5 independent biological repeats. Comparisons were performed with a Student’s t test whose values are denoted in the figures as *p ≤ 0.05, **p ≤ 0.01, ***p ≤ 0.001, ****p ≤ 0.0001 and n.s.= no significance. **d,** Amount of cell death assessed by percentage of cell debris by flow cytometry. MLL-AF9/NRas^G12D^ leukemic cells were treated with Smac-mimetic birinapant (500 nM) ± tankyrase inhibitor IWR-1 (250nM, 500nM, 1 μM, 2 μM, 5 μM) for 15 hours or Smac-mimetic birinapant (20 nM) + caspase inhibitor IDN-6556 (5 μM) ± IWR-1 (250nM, 500nM, 1 μM, 2 μM, 5 μM) for 7 hours. Graphs show mean ± SEM, n=3 biologically independent repeats. Comparisons were performed with a Student’s t test whose values are denoted in the figures as *p ≤ 0.05, **p ≤ 0.01 and ***p ≤ 0.001. **e,** TNF-induced complex 2 immunoprecipitation using anti-FLAG M2 affinity beads. Western blot analysis of complex 2 and lysates from *Casp8*^C3FLAG/C3FLAG^ MEFs using the indicated antibodies is shown. Cells were treated with TNF (100 ng/mL) + Smac-mimetic (50 nM) + caspase inhibitor (5 μM) (TSI) ± tankyrase inhibitor IWR-1 (5 μM) for 2 hours before being subjected to anti-FLAG immunoprecipitation. Filled arrowheads alone indicate potential tankyrase species. Blots are representative of two to three independent experiments. **f,** Cell death monitored by time-lapse imaging of PI staining over 16 hours using IncuCyte. *Ripk1*^D325A/+^ heterozygote MDFs were treated with TNF (50 ng/mL) + Smac-mimetic (10 nM or 100nM) (TS) ± tankyrase inhibitor IWR-1 (250nM, 500nM, 1 μM, 2 μM) for 16 hours. Dashed lines denote the PI count without IWR-1 treatment for reference. The results from two biologically independent MDF lines are shown.

**Extended Data Fig. 4.**
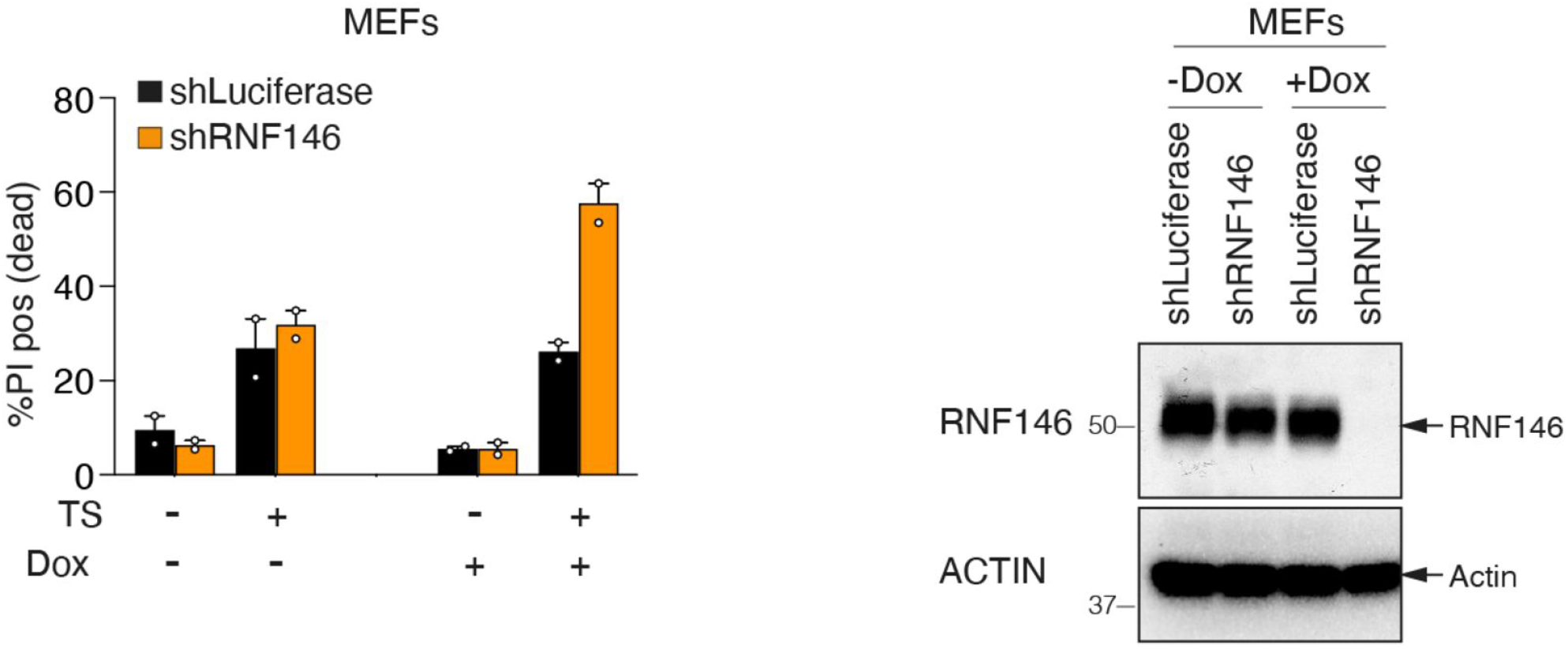
The tankyrase-RNF146 axis regulates the stability of complex 2 and TNF-induced death. Level of cell death assessed by PI positive cells. WT MEFs expressing Dox-inducible shLuciferase or shRNF146 were pre-treated with ± Dox (1 μg/mL) for 48 hours. Cells were then subjected to Western blot analysis or treated with TNF (100 ng/mL) + Smac-mimetic (25 nM) (TS) ± Dox (1 μg/mL) for another 12 hours. Blots are representative of two independent experiments. Graphs show mean ± SD throughout, n = 2 independent biological repeats.

## Acknowledgement

This work was funded by NHMRC grants 1145888, 1163581 and fellowship 1107149 (JS 2016-2020) and was made possible through Victorian State Government Operational Infrastructure Support and Australian Government NHMRC IRIISS (9000433). Research in the laboratory of M.O.H is funded by the Kanton of Zurich and the Swiss National Science Foundation (grant 310030A_176177). The generation of the *Casp8*^N3FLAG^, *Casp8*^C3FLAG^ and *Tnks2^-/-^* mice used in this study was supported by the Australian Phenomics Network (APN) and the Australian Government through the National Collaborative Research Infrastructure Strategy (NCRIS) program.

## Author contributions

L. L., J.J.S., N.L. and J.S. designed and performed experiments and interpreted data. D.M.L.P., M. O.H., J.J.S. and A.I.W. contributed reagents, analysis and interpretation. N.S., Z.Q.H., E.M., D.C., and T.K. performed experiments. A.J.K. generated the CRISPR mice. L.L., N.L. and J.S. conceived the project and wrote the paper with input from all authors.

## Materials and methods

### Mice

All mouse studies complied with relevant ethical regulations and were approved by the Walter and Eliza Hall Institute Animal Ethics Committee. The *Casp8*^N3FLAG^, *Casp8*^C3FLAG^ and *Tnks2*^-/-^ mice were generated by the MAGEC laboratory (WEHI, Australia) on a C57BL/6J background using CRISPR/Cas9. To generate *Casp8*^N3FLAG^ mice, 20 ng/μL of Cas9 mRNA, 10 ng/μL of sgRNA (CTTCTACCTCTTGATAAGAA) and 40ng/μL of the oligo donor (gatcattagcatcttgtgttgacccagGTTACAGCTCTTCTACCTCTTGATAAGAATGAATGACTACAAGGACCACGACGGTGACTACAAGGACCACGACATCGACTACAAAGACGATGACGACAAGGATTTCCAGAGTTGTCTTTATGCTATTGCTGAAGAACTGGGCAGTGAAGACCTGGCTG CCC) (in which uppercase bases denote exons; lowercase bases denote intron sequences) were injected into the cytoplasm of fertilized one-cell stage embryos generated from wild-type (WT) C57BL/6J breeders. To generate *Casp8*^C3FLAG^ mice, 20 ng/μL of Cas9 mRNA, 10 ng/μL of sgRNA (CCAGGAGGCCAAACTTACTG) and 40ng/μL of the oligo donor (GATCCTGTGAATGGAACCTGGTATATTCAGTCACTTTGCCAGAGCCTGAGGGAAAG ATGTCCTCAAGGAGATGACATTCTTAGCATCCTGACTGGCGTGAACTATGACGTGAGC AATAAAGACGACAGGAGGAACAAGGGAAAGCAGATGCCACAGCCCACCTTCACACTA CGGAAGAAGCTCTTCTTCCCTCCCGACTACAAGGACCACGACGGTGACTACAAGGACC ACGACATCGACTACAAAGACGATGACGACAAGtaatgaAGgtaagtttggcctcctgggcccctctcagggtt atgcttccttactcatttctgtggtta) were injected into the cytoplasm of fertilized one-cell stage embryos generated from WT C57BL/6J breeders. To generate *Tnks2^-/-^* mice, 20 ng/μL of Cas9 mRNA, 10 ng/μL of sgRNA (CTACACTACACCCGTATGGC and GGTTCCCCTCATTCAGACGC) were injected into the cytoplasm of fertilized one-cell stage embryos generated from WT C57BL/6J breeders. 24 hours later, two-cell stage embryos were transferred into the uteri of pseudo-pregnant female mice. Viable offspring were genotyped by next-generation sequencing. Targeted animals were backcrossed twice to WT C57BL/6J to eliminate off-target mutations.

### Cells

BMDMs were isolated from the tibia and femur of mice. MEFs were isolated from E14 embryos and MDFs were isolated from mouse tails. After SV40 transformation, MEFs and MDFs were tested for mycoplasma. 293T cells (ATCC) were used to produce SV40 viruses. HT29 were purchased from ATCC. HT1080 were gifts from Prof. John Mariadason. *Casp8^-/-^.Mlkl^-/-^* MDFs and *Mlkl^-/-^* MDFs were generated by Dr Maria Tanzer (WEHI, Australia). *Ripk1*^D325A^ MDFs were generated by Dr Najoua Lalaoui (WEHI, Australia). MLL-AF9/NRas^G12D^ leukemic cells were generated by Dr Gabriela Brumatti (WEHI, Australia).

### Reagents

The Smac-mimetic compound A (Comp A), birinapant, the caspase inhibitor IDN-6556 (Idun Pharmaceuticals) and the RIPK1 inhibitor necrostatin-1 were synthesized by TetraLogic Pharmaceuticals. Recombinant Fc-TNF was produced in house. Lyophilised human TNF was a kind gift from Prof. Dr. Daniela N. Männel. MG132 (M7449), doxycycline (D9891), cycloheximide (C4859) and the tankyrase inhibitor IWR-1 (I0161) and the deubiquitinating enzyme inhibitor N-Ethylmaleimide (NEM) (E3876) were from Sigma. The tankyrase inhibitor Az6102 (S7767) and PARP1/2 inhibitor olaparib (S1060) were from Selleckchem. PARG was generated in house by M.O.H (University of Zurich). The PARG inhibitor ADP-HPD was from Enzo (ALX-480-094-C060). 3x FLAG peptide was from Apex Bio (A6001).

### Plasmids

Constructs were designed by J.S. and synthesized by Genscript (Nanjing, CN) except for N-, C-3x FLAG tagged and tagless murine caspase-8 constructs (in house). In brief, inserts were generated by polymerase chain reaction (PCR) and fragments were sub-cloned into the pFTRE3G vector backbone. Fragments and vectors were ligated using *Bam*HI *Nhe* I sites. Restriction enzymes, T4 DNA ligase and corresponding buffers were used as per manufacturer’s instructions ^93^. Ligation products were transformed into XL1-Blue competent cells (Agilent Technologies) and constructs were purified by miniprep kit (QIAGEN). Construct sequences were verified by Sanger sequencing performed by the Australian Genome Research Facility (AGRF).

### Inducible shRNA generation

The Dox-inducible pF H1tUTG-GFP shRNA vector was kindly provided by A/Prof. Marco Herold (WEHI, Australia). The sequences of shRNAs are listed as following:

shLuciferase sense: 5’-tcccTGCGTTGCTAGTACCAACttcaagagaGTTGGTACTAGCAACGCA tttttc-3’

shLuciferase antisense: 5 ‘-tcgagaaaaaTGCGTTGCTAGTACCAACtctcttgaaGTTGGTACTAGCA ACGCA-3’

shScrambled sense: 5’-tcccTTCTCCGAACGTGTCACGTttcaagaga ACGTGACACGTTCGGAG AAtttttc-3’

shScrambled antisense: 5’-tcgagaaaaaTTCTCCGAACGTGTCACGTtctcttgaa ACGTGACACGTT CGGAGAA-3’

shRNF146 sense: 5’-tcccATTTCTGCCCACGTAACATTAttcaagagaTAATGTTACGTGGGCA GAAATtttttc-3’

shRNF146 antisense: 5’-tcgagaaaaaATTTCTGCCCACGTAACATTAtctcttgaaTAATGTTACGTGGGCAGAAAT-3’

shTNKS 1-1 sense: 5’-tcccCGTCTCTTAGAGGCATCGAAAttcaagagaTTTCGATGCCTCTAAGAGACGtttttc-3’

shTNKS 1-1 antisense: 5’-tcgagaaaaaCGTCTCTTAGAGGCATCGAAAtctcttgaaTTTCGATGCCTCTAAGAGACG-3’

shTNKS 1-2 sense: 5’-tcccGCTCCAGAAGATAAAGAATATttcaagagaATATTCTTTATCTTCTGGAGCtttttc-3’

shTNKS 1-2 antisense: 5’-tcgagaaaaaGCTCCAGAAGATAAAGAATATtctcttgaaATATTCTTT ATCTTCTGGAGC-3’

shRNAs and pF H1tUTG-GFP vectors were ligated following *Xho* I */Bsm* BI restriction digestion. shRNA cell lines were generated by infecting indicated cells with lentivirus containing Dox-inducible control shRNAs or shRNA targeting murine TNKS1 or RNF146 followed by fluorescence-activated cell sorting (FACS) for GFP fluorescent signal (excitation/emission= 488/509 nm).

### Immunoprecipitation

For complex 1 purification, MEFs were seeded in 15 cm dishes and treated as indicated with Fc-TNF (1 μg/mL). Cells were lysed in DISC lysis buffer (150 mM sodium chloride, 2 mM EDTA, 1% Triton X-100, 10% glycerol, 20 mM Tris, pH 7.5). Protein lysates were immunoprecipitated with protein A Sepharose (40 μL/sample, WEHI antibody facility, Australia) for 4 hours at 4°C. Beads were washed 4 times with DISC and samples were eluted by boiling in 1x SDS loading dye. For complex 2 purification, cells were seeded in 15 cm dishes and treated as indicated. Cells were lysed in DISC. Protein G or protein A Sepharose (20 μL/sample, WEHI antibody facility, Australia) pre-blocked with DISC lysis buffer containing 2% BSA were bound with indicated antibody (1.5 μg antibody/ sample). Anti-cleaved caspase-8 (4790) and anti-RIPK1 (3493) were from cell signalling technology (CST). Anti-FADD (clone 7A2) was produced in house. Anti-tankyrase antibody (sc-365897) was from Santa Cruz Biotechnology. Anti-PAR (4335-MC-100) was from Trevigen. Protein lysates were precipitated at 4 °C overnight. Beads were washed 4 times with DISC and samples were eluted by boiling in 1x SDS loading buffer. For anti-FLAG immunoprecipitation, ANTI-FLAG® M2 Affinity Gel (15 μL/sample, Sigma) were blocked with DISC lysis buffer containing 2% BSA for 1 hour at 4°C and incubated with protein lysates at 4 °C overnight. After washing 4 times with DISC, samples were eluted with FLAG peptides (1mg/mL) and denatured by boiling in 5 x SDS loading buffer at 100°C for 15min.

### GST pulldown assay

For enrichment of PARylated proteins, plasmids pGEX 6P3 hs GST RNF146 WWE, pGEX 6P3 hs GST RNF146 WWE R163A, pGEX 6P3 hs HUWE1 WWE, pGEX 6P3 mm TRIP12 WWE, pGEX 6P3 AF1521 ^49,50,56^ were designed by J.S. and synthesized by Genscript (Nanjing, CN). Plasmids were transformed into BL21 *E. coli* (DE3) (Thermo Fisher) and grown at 37°C to an optical density (600 nm) of ~0.6-0.8 in Super Broth before protein expression was induced with 1 mM IPTG (Sigma) overnight at 18°C. Recombinant protein was purified by Glutathione Xpure Agarose Resin (UBPBio) and size exclusion chromatography (SEC). BMDMs, MEFs or MDFs were treated as indicated and cells were lysed in DISC lysis buffer supplemented with 5 μM ADP-HPD. Equal protein amounts were incubated with Glutathione Sepharose® 4B (GE Healthcare) charged with GST fusion proteins overnight at 4°C. After washing 4 times with PBST buffer (PBS + 0.2% Tween-20 (Sigma)), samples were eluted by boiling at 100°C in 1x SDS loading buffer for 15min. For enrichment of ubiquitylated proteins, MEFs or MDFs were treated as indicated and cells were lysed in DISC lysis buffer supplemented with 10 mM NEM and incubated with Glutathione Sepharose^®^ 4B pre-coupled with GST-UBA1 fusion protein (produced by Aleksandra Bankovacki, WEHI, Australia) overnight at 4°C. After washing 4 times with PBST buffer, samples were eluted by boiling in 1x SDS loading buffer at 100°C for 15min.

### PARG treatment

Recombinant PARG generated in house by M.O.H. Immunoprecipitated TNFR1 complex 2 was eluted with FLAG peptides and the immunoprecipitants were diluted with 2x PARG reaction buffer (100 mM KH2PO4, 100 mM KCl, 0.2 mg/mL BSA, 0.2% Triton X-100). Recombinant PARG (7.2 μg/sample) was added and the reaction mixture was incubated at 37°C for 3 hours, followed by GST-WWE pulldown overnight at 4°C.

### Western blotting

Cells lysates were separated on 4-2% gradient SDS-polyacrylamide gels (Biorad), transferred to polyvinylidene fluoride (Millipore) membranes and blotted with indicated antibodies purchased from CST except for caspase-8 (M058-3, MBL Life Science), RIPK3 (33/16-8G7-1-1, WEHI antibody facility, Australia), phospho-RIPK3 (a gift from Genentech), phospho-MLKL (ab196436, Abcam), FADD (ADI-AAM-212-E, Enzo), tankyrases (sc-365897, Santa Cruz Biotechnology), PAR (MABC547, Sigma), RNF146 (73-233, NeuroMab), cIAP1 (clone 1E1-1-12, in house), actin (A1978, Sigma) and HSP90 (ADI-SPA-835, Enzo).

### Flow cytometry

2 x 10^4^ MEFs or MDFs were seeded in 96 well plates and 2 x 10^5^ BMDMs were seeded in 48 well plates. 24 hours later cells were treated as indicated for the indicated times. Cells were then trypsinized and collected into 1.2 mL FACS tubes. Propidium iodide (PI, 1 mg/mL) was added and cells were spun down at 300 g for 5 min at 4°C. PI signal was excited by Blue laser (488 nM) and the emission was received through B660LP filter. 1×10^4^ of MLL-AF9/NRas^G12D^ cells were seeded in 96 well plates and treated as indicated for the indicated times. Cell death was analysed by flow cytometry using a FACSCalibur (BD Biosciences).

### Time-lapse imaging (IncuCyte)

Percentage cell death was assayed every 45 min to 1 hour by time-lapse imaging using the IncuCyte live cell analysis imaging (Essenbioscience) for 16-24 hours with 5% CO2 and 37°C climate control. Dead cells were identified by PI (0.25 μg/mL) staining. PI was added to the cells 2 hours before imaging and compounds were added 10 min before the start of imaging. Dead cells were counted using the in-built ‘Basic’ analysis software and PI positive cells were calculated based on the imaged area.

### Mass spectrometry

After anti-FLAG immunoprecipitation, samples were eluted with FLAG peptide and then added to a filter-aided sample preparation (FASP) column (J.J.S., WEHI, Australia) and spun (14,000 g) until volume hadpassed through the column. Protein material was reduced with tris(2-carboxyethyl) phosphine (TCEP; 10 mM final) and digested overnight with 2 μg sequence-grade modified trypsin Gold (Promega, V5280) in 50 mM ammonium bicarbonate (NH4HCO3) at 37°C. Peptides were collected into microvial tubes and acidified with formic acid (FA) to a final concentration of 1% (v/v). Samples were frozen at −80°C and subsequently lyophilised.

Peptides were resuspended in 2% (v/v) acetonitrile (ACN) and 1% (v/v) FA and injected and separated by reversed phase liquid chromatography on a M-class UHPLC system (Waters, USA) using a 250 mm x 75 mm column (1.7mm C18, packed emitter tip, Ion Opticks, Australia) with a linear 90 min gradient at a flow rate of 400 nl/min from 98% (v/v) solvent A (0.1% (v/v) FA in Milli-Q water) to 35% (v/v) solvent B (0.1% (v/v) FA, 99.9% (v/v) ACN). The nano-UHPLC was coupled on-line to a Q-Exactive Orbitrap mass spectrometer equipped with an EASY-spray ionization source (Thermo Fisher Scientific, Germany). The Q-Exactive was operated in a data-dependent mode, switching automatically between one full-scan and subsequent MS/MS scans of the ten most abundant peaks. The instrument was controlled using Exactive series version 2.8 build 2806 and Xcalibur 4.0. Full-scans (m/z 350–1,850) were acquired with a resolution of 70,000 at 200 m/z. The 10 most intense ions were sequentially isolated with a target value of 1e5 ions and an isolation width of 2 m/z and fragmented using higher-energy collisional dissociation with stepped normalised collision energy of 19.5, 26, 32. Maximum ion accumulation times were set to 80 ms for full MS scan and 200 ms for MS/MS.

All raw files were analyzed by MaxQuant v1.6.10.43 software using the integrated Andromeda search engine. Data was searched against the mouse Uniprot Reference Proteome with isoforms (downloaded March 2018) and a separate reverse decoy database using a strict trypsin specificity allowing up to 2 missed cleavages. The minimum required peptide length was set to 7 amino acids. Modifications: Carbamidomethylation of Cys was set as a fixed modification, while N-terminal acetylation of proteins and oxidation of Met were set as variable modifications. First search peptide tolerance was set at 20 ppm and main search set at 4.5 ppm (other settings left as default). Matching between runs and LFQ quantitation was turned on. Maximum peptide mass [Da] was set at 8000. All other settings in group or global parameters were left as default.

Further analysis was performed using a custom pipeline developed in R (3.6.1), which utilizes the LFQ intensity values in the MaxQuant output file proteinGroups.txt. Proteins not found in at least 50% of the replicates in one group were removed. Missing values were imputed using a random normal distribution of values with the mean set at mean of the real distribution of values minus 1.8 s.d., and a s.d. of 0.3 times the s.d. of the distribution of the measured intensities. The log2 fold changes and probability of differential expression between groups was calculated using the Limma R package (3.4.2). Probability values were corrected for multiple testing using Benjamini-Hochberg method.

### Statistical analyses

The number of independent experiments for each dataset is stipulated in the respective figure legend. Comparisons were performed with a Student’s t test whose values are represented in the figures as *p ≤ 0.05, **p ≤ 0.01, ***p ≤ 0.001 and ****p ≤ 0.0001 and n.s.= no significance using Prism v.8.2 (GraphPad).

